# Natural Music Evokes Correlated EEG Responses Reflecting Temporal Structure and Beat

**DOI:** 10.1101/705517

**Authors:** Blair Kaneshiro, Duc T. Nguyen, Anthony M. Norcia, Jacek. P. Dmochowski, Jonathan Berger

## Abstract

The brain activity of multiple subjects has been shown to synchronize during salient moments of natural stimuli, suggesting that correlation of neural responses indexes a brain state operationally termed ‘engagement’. While past electroencephalography (EEG) studies have considered both auditory and visual stimuli, the extent to which these results generalize to music—a temporally structured stimulus for which the brain has evolved specialized circuitry—is less understood. Here we investigated neural correlation during natural music listening by recording dense-array EEG responses from N = 48 adult listeners as they heard real-world musical works, some of which were temporally disrupted through shuffling of short-term segments (measures), reversal, or randomization of phase spectra. We measured neural correlation across responses (inter-subject correlation) and between responses and stimulus envelope fluctuations (stimulus-response correlation) in the time and frequency domains. Stimuli retaining basic musical features evoked significantly correlated neural responses in all analyses. However, while unedited songs were self-reported as most pleasant, time-domain correlations were highest during measure-shuffled versions. Frequency-domain measures of correlation (coherence) peaked at frequencies related to the musical beat, although the magnitudes of these spectral peaks did not explain the observed temporal correlations. Our findings show that natural music evokes significant inter-subject and stimulus-response correlations, and suggest that the neural correlates of musical engagement may be distinct from those of enjoyment.

## Introduction

Over the past 15 years, neuroscience has begun to unravel the brain’s processing of natural audiovisual stimuli. These advances have been facilitated by the discovery that neural responses are significantly correlated across individuals or stimulus repetitions (Hasson et al., 2010). This inter-subject correlation (ISC) approach originated in functional magnetic resonance imaging (fMRI) research (Hasson et al., 2004) and has more recently gained traction with electroencephalography (EEG). Importantly, ISC of EEG responses to natural stimuli has been shown to reflect numerous cognitive and behavioral states, including memory retention (Cohen & Parra, 2016), top-down attention (Ki et al., 2016), perception of time (Cohen et al., 2017), audience preferences (Dmochowski et al., 2014), and level of consciousness (Iot-zov et al., 2017). The purported mechanism underlying these effects is the ‘synchronization’ of individual neural responses by a common audiovisual stimulus, leading to elevated levels of ISC during moments marked by increased affect, attention, and memory recall—a brain state that has been defined by some as ‘engagement’(Hasson et al., 2004; Dmochowski et al., 2012; Lankinen et al., 2014). In a related approach termed stimulus-response correlation (SRC), time-varying stimulus features are correlated with the evoked EEG responses. SRC has been shown to covary with ISC and to reflect attentional states of participants (Dmochowski et al., 2018). These approaches have potential widespread utility in both understanding brain activity in real-world settings and developing technologies to decode brain state from measures of neural activity.

To date, investigations of EEG correlation with natural stimuli have primarily considered narrative works in audiovisual (Dmochowski et al., 2012, 2014; Cohen & Parra, 2016;Ki et al., 2016; Cohen et al., 2017; Poulsen et al., 2017; Dmochowski et al., 2018) and audio (Cohen & Parra, 2016; Ki et al., 2016; Iotzov et al., 2017) modalities. Given that methods for studying the brain’s processing of natural stimuli are indispensable, it is critical to investigate the extent to which the correlation approach can be applied to responses to natural music. Instrumental music, particularly from the Western classical tradition of the 18–19th centuries, is often compared to narrative (Maus, 1991). However, a number of factors challenge the notion of music as a narrative art and, for the purposes of this study, distinguish music for separate consideration.

Correlation of neural responses to natural music has been performed in various imaging modalities. fMRI studies have linked neural correlation during music listening to computationally extracted, time-varying acoustical features (Alluri et al., 2012; Trost et al.,2015) and have shown that perturbing music’s temporal and spectral characteristics reduces ISC (Abrams et al., 2013). In a recent fMRI study, Farbood et al. (2015) computed ISC of responses to intact music, and to control stimuli scrambled at perceptually relevant timescales such as measures and phrases, identifying a hierarchy of brain areas corresponding to different levels of music’s structural temporal hierarchy. An ISC approach has been applied to electrocorticographic (ECoG) responses to natural music to identify task-related cortical regions and spatial relationships between them (Potes et al., 2014). Most recently, EEG studies have identified optimal stimulus and response filters to maximize SRC (Gang et al., 2017) and have assessed the impact of familiarity, repeated listens, and musical training on ISC (Madsen et al., 2019) during natural music listening.

In a separate line of research, EEG responses to rhythmic auditory stimuli have been analyzed using a ‘frequency tagging’ approach. This approach leverages the finding that periodic stimuli elicit steady-state evoked potentials (SS-EPs) with spectral peaks at frequencies related to stimulus periodicities (see Norcia et al. (2015) for review). While the SS-EP is most often studied in response to visual stimuli, periodicities in the EEG have also been observed in auditory studies involving controlled, synthesized rhythmic stimuli, implicating frequencies related to musical beat and meter (e.g., Nozaradan et al. (2011); Nozaradan (2014); Lenc et al. (2018)). However, the extent to which stimulus-related periodicities are reflected in responses to complex, natural music is less known, particularly in the case of long, continuous response epochs.

In the present study we conducted an EEG investigation of neural correlation during natural music listening. We hypothesized that ISC, SRC, and listener enjoyment would be highest for musical excerpts that retained the temporal organization with which they were composed. To test this hypothesis, we recorded scalp EEG from N = 48 human participants as they listened to Hindi-language (‘Bollywood’) songs that have attained widespread commercial success but were new to our listeners. Importantly, in some cases the songs had been edited to disrupt the natural musical ordering of acoustic events (for example, played in reverse or with the measures scrambled in time), and we expected to observe reduced enjoyment and neural correlation for these excerpts. By utilizing stimuli comprising temporal regularities through a steady beat, ISC and SRC could be investigated in the frequency domain as well, laying groundwork for a coherence approach—a measure of neural correlation that music can uniquely provide. To the best of our knowledge, this is the first EEG correlation study to consider temporal manipulations of natural music stimuli; to study music-evoked ISC and SRC in tandem; and to employ the coherence approach in order to identify correlated frequencies in responses to natural music.

## Methods

### Stimuli

Stimuli were derived from four songs that appeared in recent Hindi-language ‘Bollywood’ films (Table 1). These songs were widely popular due to their inclusion in top-grossing films (Rao, 2007). Therefore, they have successfully engaged a massive audience but would be new to our participant pool, avoiding possible confounds of familiarity and established preference. Moreover, the songs—falling broadly in the pop genre with verse and chorus elements, clear vocal lines, easily discernible phrase structure, and arguably Western instrumental, melodic, and timbral palettes—were composed to be easy to grasp and enjoy upon first listen. Lyrics were sung in Hindi and Hindi dialects, containing at most sporadic, single-word English utterances; thus, they would not be comprehensible to our participant pool. Consequently, the listening experience would not be impacted by content of the lyrics, and stimulus manipulations would bring about no loss of lyrics-related meaning. Finally, our planned stimulus manipulations required songs with a steady beat throughout, which was easy to achieve in this genre.

**Table 1:** Song title, film title, year of film release, length (min:sec), and tempo (BPM) of the four songs forming the basis of the stimulus set.

|  | **Song 1** | **Song 2** | **Song 3** | **Song 4** |
| --- | --- | --- | --- | --- |
| **Title** | Ainvayi Ainvayi | Daaru Desi | Haule Haule | Malang |
| **Film** | *Band Baaja Baaraat* | *Cocktail* | *Rab Ne Bana Di Jodi* | *Dhoom 3* |
| **Year** | 2010 | 2012 | 2008 | 2013 |
| **Length** | 4:27 | 4:30 | 4:24 | 4:33 |
| **Tempo** | 156 | 94 | 90 | 86 |

We purchased digital versions of the songs from iTunes and used Audacity software^1^ to convert the files to .wav format. Further processing of the stimuli was performed in Matlab.^2^ Each audio file was converted from stereo to mono by computing the mean of the channels. Next, we used publicly available beat-tracking software (Ellis, 2007) to compute beat onset times, which were manually corrected to account for tempo octave errors (Levy, 2011;Moelants & McKinney, 2004) as verified by four trained musicians. Using corresponding measure onsets—which for the selected songs denoted hierarchical groupings of four beats—we trimmed excess silence from the beginning and end of each audio file so that every song comprised an integer number of measures. We henceforth refer to the resulting audio files as the Intact versions of the songs.

The experiment comprised 16 stimuli: Four songs each in Intact, Measure, Reversed, and Phase conditions (see Figure 1A for Song 1 waveforms). All stimuli were around 4 minutes and 30 seconds in length and were presented in their entirety. In addition to the Intact stimuli described above, we devised three stimulus manipulations to impose varying degrees of temporal disruption while preserving each song’s aggregate frequency content.

**Figure 1:**
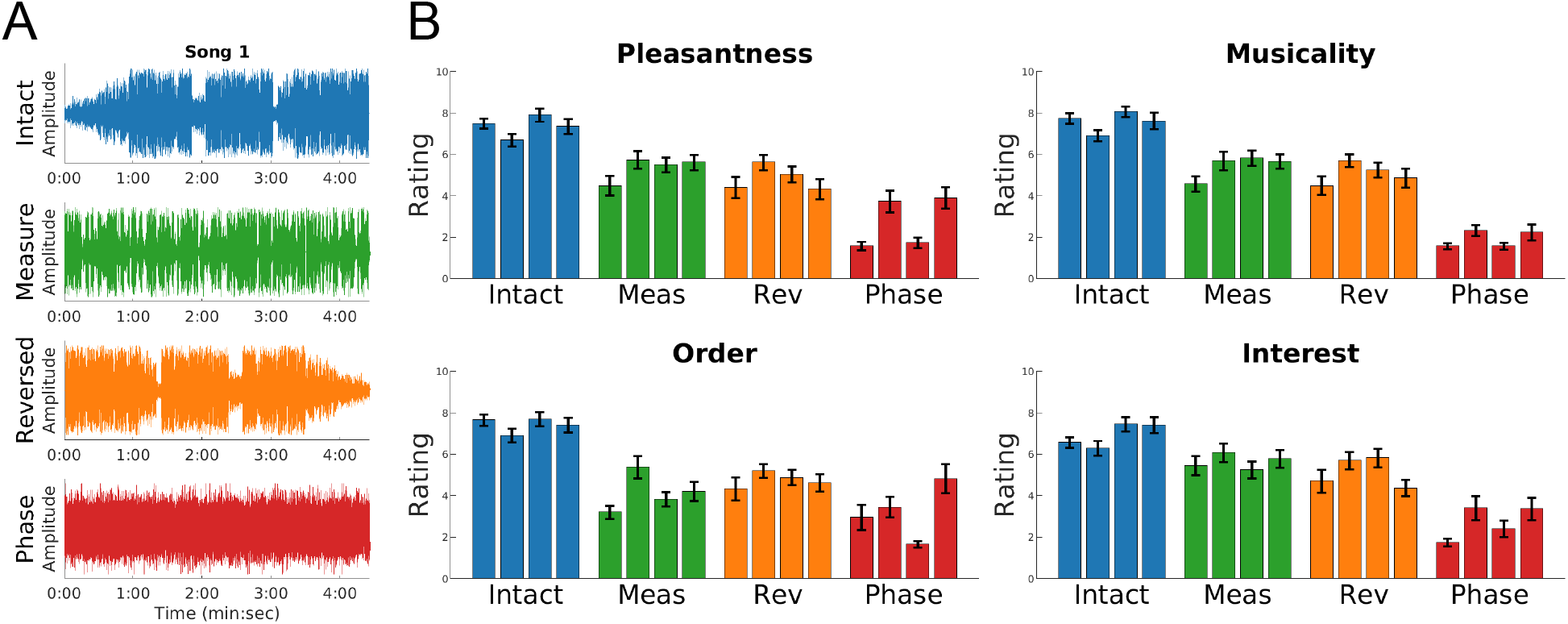
Intact songs received highest behavioral ratings. (**A**) Song 1 stimulus waveforms: Original (Intact-blue), shuffled at the measure level (Measure-green), played in reverse (Reversed-orange), and with the phase spectrum randomized (Phase-red). (**B**) Behavioral ratings of Pleasantness, Musicality, Order, and Interest varied significantly by stimulus condition (repeated-measures ANOVA, *N* = 48 participants, *p* < 0.001, FDR corrected). Intact stimuli received highest ratings (all *p* < 0.001, FDR corrected) and Phase stimuli received lowest ratings (all *p* < 0.01, FDR corrected). Error bars denote twice the standard error of the mean (*N* = 24 trials per stimulus).

The first manipulation was measure shuffling (Measure), achieved by re-computing beat onset and measure boundary times of Intact audio files, and then partitioning and randomly shuffling each song at its measure boundaries. The resulting audio preserved short-term temporal organization as well as such features as beat, meter, tempo, and instrumentation while disrupting longer-term musical trajectories such as phrases and song parts. This approach is analogous to the scene-scrambling manipulation used by Dmochowski et al. (2012) and was used in the recent music fMRI ISC study by Farbood et al. (2015). The second manipulation was time reversal (Reversed)—an approach also used by Farbood et al. (2015), and prior to that with narrative visual and auditory stimuli (Hasson et al., 2008; Regev et al., 2013). In the case of music, time reversal disrupted tonal and structural trajectories and in some cases timbral characteristics of the songs while preserving instantaneous sensory events as well as beat, meter, tempo, and phrase- and song-part-level grouping. The final and most extreme manipulation was achieved by phase scrambling the stimuli (Phase). We transformed each Intact waveform to the frequency domain and independently randomized the phase of each positive frequency to a value between 0 and 2*π*. Following this, we assigned conjugate-symmetric phases to negative frequencies and transformed the signal back to the time domain (Prichard & Theiler, 1994). This procedure left aggregate magnitude spectra unchanged while disrupting all temporal structure, resulting in what sounds like shaped noise. This approach was used to produce a control version of a symphonic work in a recent fMRI study by Abrams et al. (2013).

Stimuli were assigned to participants according to a planned Latin square design, whereby each participant would be assigned each song once, with a different stimulus condition for each song. This procedure guaranteed that a participant heard each song in only one condition, and therefore had no basis for comparing different conditions of a song. A total of 4! = 24 such assignments were possible; we planned to use each assignment twice across a pool of 48 participants. In all, 12 participants would be assigned to each of the 16 stimuli.

As a stimulus feature we used magnitude envelope fluctuations. Each stimulus envelope was computed as the absolute value of the audio waveform’s Hilbert transform, and then resampled to 125 Hz to match the sampling rate of the EEG. Envelope fluctuations were computed as the absolute value of the temporal derivative of the z-scored envelope.

### Participants

Data from *N* = 48 right-handed adults (25 males), 18–34 years old (mean 24.58 years) with normal hearing were analyzed in this study. In order to obtain 48 complete, usable datasets, it was necessary to collect data from 58 participants. Unusable datasets were flagged during data collection and preprocessing—prior to any spatial filtering and correlation analyses. Reasons for exclusion were gross noise artifacts during data collection (*N* = 3), 20 or more bad electrodes identified during preprocessing (*N* = 4), participant did not follow instructions during the experimental session (listening with eyes closed, *N* = 1), and participant revealed during data collection that they did not meet eligibility criteria for the experiment (*N* = 2). When a participant was excluded, their stimuli and stimulus orderings were re-assigned to a subsequent participant.

All participants were fluent in English and had no cognitive or decisional impairments. We recruited participants who reported enjoying music from various genres including pop, rock, and classical, and who listened to music for at least three hours per week. Participants were additionally required to have no experience with Hindi language or media including films and music. As we wished for our experimental findings to generalize across varying levels of musical expertise, we had no requirements related to formal musical training or ability except that participants not have absolute pitch, as this is known to impact cortical organization for music processing (Loui et al., 2011). Thirty-two participants reported formal training, ranging from three months to 22 years (mean 7.57 years). Twenty participants reported being currently involved in musical activities. Reported music listening ranged from 3–52.5 hours (mean 15.03 hours) per week.

### Experimental Paradigm and Data Acquisition

This research was approved by the Stanford University Institutional Review Board. All participants delivered written informed consent prior to completing any experimental interventions. After each participant delivered informed consent, he or she filled out a demographic and musical experience questionnaire. Following this, the participant completed a training block containing two 15-second stimuli not used in the actual experiment in order to be familiarized with the task and procedure for delivering behavioral ratings. Once the participant was comfortable with the experimental paradigm, he or she was fitted with an electrode net and completed the experiment, which consisted of two recording blocks of around 20 minutes each. During each block, the participant heard each assigned stimulus once. Participants were instructed to listen attentively and watch a fixation cross presented on a computer monitor while stimuli were playing. At the conclusion of each stimulus, the participant used a computer keyboard to rate the excerpt, on a scale of 1–9, along dimensions of Pleasantness, Musicality, Order, and degree of Interest. Stimulus ordering was randomized for every participant and experimental block. As each participant heard their assigned stimuli twice and 12 participants were assigned to each stimulus, we collected a total of 24 trials for each of the 16 stimuli.

The experiment was programmed using Neurobehavioral Systems Presentation software.^3^ The stimulus presentation computer was synced to a monitor located 57 cm in front of the participant, as well as a keyboard and mouse, in an acoustically and electrically shielded ETS-Lindgren booth. Stimuli were delivered via two magnetically shielded Genelec 1030A speakers located 120 cm from the participant in the booth. Stimulus labels and participant key press responses were registered by the Presentation software and sent to the EEG amplifier. Precise time-stamping of stimulus onsets was achieved by sending intermittent square-wave pulses to the EEG amplifier directly from the second audio channel of the stimulus, which was not played to participants.

Data were acquired using the Electrical Geodesics, Inc. (EGI) GES 300 platform (Tucker, 1993). EEG was recorded using a Net Amps 300 amplifier and unshielded 128-channel HCGSN 110 and 130 nets at a sampling rate of 1 kHz with vertex reference. Electrode impedances were verified to be below a threshold of 60 kΩ prior to the start of each recording (Ferree et al., 2001). Amplified EEG was recorded using Net Station software.

### Data Preprocessing

EEG recordings were filtered using the built-in Net Station Waveform Tools zero-phase bandpass filter (0.3–50 Hz), and temporally downsampled by a factor of 8 to a sampling rate of 125 Hz. Each recording was then exported to Matlab .mat file format. All subsequent analyses were performed using in-house Matlab code unless otherwise specified.

Preprocessing was performed on a per-recording basis as follows. We first extracted the behavioral ratings delivered by the participant at the end of each trial. Next, the continuous EEG was epoched using the time stamps sent from the audio and associated with corresponding stimulus labels sent from Presentation. As we observed zero-frequency content in the recordings even after EGI filtering, we removed any linear trend and performed median-based DC correction on a per-trial basis. Trial epochs were then concatenated in time. We computed electrooculogram (EOG) channels using electrodes above and below the eyes (vertical) and at the right and left outer canthi (horizontal), retained electrodes 1–124 for further analysis (excluding electrodes on the face), and excluded electrodes flagged as bad during data collection. Ocular artifacts were removed semi-automatically using Independent Components Analysis (Bell & Sejnowski, 1995; Jung et al., 1998) using the extended Infomax `runICA` implementation from the Matlab EEGLAB toolbox (Delorme & Makeig, 2004). Independent components correlating with either EOG component at |*r*| ≥ 0.3 were automatically replaced with zeros; additional components with magnitude EOG correlation 0.2 ≤ |*r*| < 0.3 were also set to zero based on manual inspection of both the temporal activations and forward-model component topographic maps (Parra et al., 2005; Jung et al., 1998). The data were then transformed from component space back to electrode space. Following this, any electrode for which at least 10% of voltage magnitudes exceeded 50 *μV* across the concatenated trials was flagged as a recording-wide bad electrode; if any such electrode was identified, the preprocessing procedure was re-started with that electrode excluded.

Once no additional recording-wide bad electrodes were identified, the following steps were performed on a per-trial basis. First, any electrode with 10% or more of voltage magnitudes exceeding 50 *μV* in a given trial was flagged as a bad electrode and removed from that trial’s data frame. Noisy transients were addressed by replacing with NaN any data samples whose magnitude voltage exceeded four standard deviations of its electrode’s mean power, with the procedure repeated iteratively four times. Next, rows representing recording- or trial-wide bad electrodes were reconstituted with rows of NaNs, ensuring that all data records contained the same number of rows. We then appended a row of zeros to the data frame to represent the vertex reference, converted the data to average reference, imputed missing values with the spatial average of data from neighboring electrodes, and performed a final DC correction. In the end, each trial of cleaned data was an electrode-by-time matrix of size 125 × *T*, where *T* varied according to the length of the stimulus. Once all EEG recordings were preprocessed, data were aggregated across trials on a per-stimulus basis, producing for each stimulus an electrode-by-time-by-trial matrix of size 125 × *T* × 24.

### Reliable Components Analysis

To mitigate problems of high data dimensionality and low signal-to-noise ratio inherent to EEG, we spatially filtered the data using Reliable Components Analysis (RCA) (Dmochowski et al., 2012, 2015). Given a collection of electrode-by-time matrices *X_i_* ∈ ℝ*^D×T^* representing responses to a common stimulus, RCA computes linear weightings of electrodes (*w* ∈ ℝ^*D*^) to maximize correlation, in time (ℝ^*T*^), of the projected data across response trials. The optimization procedure, described extensively in Dmochowski et al. (2012, 2015), reduces to an eigenvalue solution whereby the resulting weight vectors *w_i_* and their coefficients are the ordered eigenvectors and eigenvalues, respectively, of across-trial covariance relative to within-trial covariance (Bookstein & Mitteroecker, 2014).

We performed RCA using a publicly available Matlab implementation.^4^ For each calculation we computed five RCs, retaining only the first seven principal components of the pooled covariance eigenvalue spectrum for regularization. RCA was computed on a per-condition basis, with responses to each stimulus arranged as described in Dmochowski et al.(2012). Each RCA calculation produced an electrode-by-component weight matrix *W*—the eigenvectors—as well as a vector *d* of corresponding eigenvalues (correlation coefficients). The matrix of forward-model projections of the weights *A*, used to visualize scalp topographies of the spatial filters, was computed from *W* and the pooled within-trial covariance matrix (Parra et al., 2005; Haufe et al., 2014). Also output by RCA were the projected EEG data, a 3D matrix of size time-by-component-by-trial for each stimulus, on which all subsequent analyses were performed.

### Time-Domain Analyses

We focused our analyses on the time courses of the maximally correlated component (RC1). We computed ISC on a per-stimulus basis by correlating RC1 time courses over the full duration of the stimulus. We used a one-against-all procedure, whereby the ISC of each trial was the mean correlation of that trial with all other trials. We report the mean ISC across all *N* = 24 trials for a given stimulus, as well as the standard error of the mean. To determine whether envelope properties of the stimuli (as opposed to the brain state of the listener) drove ISC, we additionally computed ISC over short (5-sec) time segments and correlated these values with the magnitude of the stimulus envelope’s temporal derivative—a measure of absolute envelope change during the segment—in the corresponding temporal windows. If abrupt changes in envelope were driving ISC, we would expect these correlations to be significantly greater than zero.

We computed SRC between each RC1 time course and the magnitude fluctuations of the corresponding stimulus envelope. As the delay between stimulus events and corresponding evoked responses is unknown, we temporally filtered each envelope feature to maximize its correlation with the already spatially filtered RC1 activations. Temporal filtering was performed separately for each stimulus across the full time course of stimulus and response. To prepare the data for this procedure, the time-by-trials RC1 EEG matrix *X* ∈ ℝ^*T*×24^ was reshaped to a vector *x*_cat_ ∈ ℝ^24*T*^ concatenating the trials in time. Next, the stimulus feature *z* ∈ ℝ^*T*^ was expanded to a Toeplitz matrix *Z* ∈ ℝ^*T*×*K*^, whose columns comprised successively sample-wise delayed versions of the envelope up to one second, plus an intercept column. This matrix was then repeated row-wise 24 times, forming *Z*_cat_ ∈ ℝ^24*T*×*K*^. The temporal filter *H* ∈ ℝ^*K*^ was computed by regressing the temporally filtered stimulus feature matrix onto the RC1 time series: 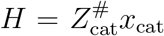, where 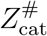 is the Moore-Penrose pseudoinverse of the stimulus Toeplitz matrix. Predicted EEG activations *x*_est_ = *Z*_cat_*H* were correlated with actual EEG responses on a per-trial basis. We report mean correlation, and standard error of the mean, across the 24 stimulus-response correlations for each stimulus.

The inter-subject and stimulus-response analyses described above produced one correlation per trial. In order to assess the relation between the two measures, we correlated per-trial ISC-SRC pairs across all stimuli (384 trials). We report the correlation coefficient and its significance.

### Frequency-Domain Analyses

Frequency-domain correlations were computed on responses to Intact, Measure, and Reversed stimuli; Phase responses were excluded because those stimuli did not have a steady beat. We computed inter-subject and stimulus-response magnitude-squared coherence on a per-stimulus basis using a DFT length of 1,024, a 5-second Hamming window, and 50% overlap between windows. This procedure output coherence magnitudes over frequencies 0–62.5 Hz (Nyquist) with a resolution of 0.122 Hz. As was done for time-domain analyses, we computed inter-subject coherence in a one-against-all fashion, while stimulus-response coherence was computed between every trial RC1 vector and the corresponding vector of stimulus envelope fluctuations (without the regression procedure). Based on visualizations of mean coherence from 0–12 Hz, we identified for each song the frequency corresponding to the most prominent coherence peak across stimulus conditions. This is henceforth termed the ‘frequency of interest’ for that song. We report the mean, and standard error of the mean, of magnitude-squared coherence at the frequency of interest for each stimulus.

The relationship between inter-subject and stimulus-response coherence was assessed by aggregating RC1 coherence measures across stimuli at each song’s frequency of interest. We then correlated the trial-wise inter-subject and stimulus-response measures. We report the correlation coefficient and its statistical significance.

We computed cross power spectral density (CPSD) phase angles using the same DFT and windowing paramaters used for coherence analyses. Inter-subject CPSD phase angles were computed on a per-trial basis using the one-against-all procedure described previously. While in the time-domain SRC analysis it was necessary to temporally filter stimulus envelopes to account for unknown delays, here the delay can be observed directly from the phase of the cross spectrum. We present the mean and standard deviation of phase angles across the 24 trials from 0–12 Hz on a per-stimulus basis. We also visualize distributions of CPSD phase at the frequency of interest of each stimulus using polar histograms.

### Statistical Analyses

The significance of each EEG analysis was assessed using the permutation testing approach described by Theiler et al. (1992). Surrogate EEG data were generated by phase-scrambling each trial matrix prior to input to RCA—in fact the same approach used to create the phase-scrambled stimuli. Phases were randomized independently for each trial, and all electrodes in a given trial were assigned the same randomized conjugate-symmetric distribution of phases. The resulting data preserved aggregate power spectra and autocorrelation characteristics inherent to EEG, but stimulus-driven temporal characteristics were lost (Sturm et al., 2014). We computed RCA over 1,000 independent instantiations of surrogate data, and the resulting component data underwent all subsequent correlation and coherence analyses. These results formed the null distribution of each analysis, against which we compared the experimental results for calculation of *p*-values.

The statistical significance of behavioral ratings (Likert ratings approximating a continuous variable, Norman (2010)), RC1 correlations, and RC1 coherence results across stimulus conditions was computed in `R` (R Core Team, 2018) using the `lme4` package (Bates et al., 2015). For each analysis, we created a linear mixed-effects model with fixed effect of stimulus condition and random effects of participant and song. As our stimulus assignment procedure did not permit within-participant paired comparisons between conditions of a given song, follow-up pairwise comparisons were performed using the same repeated-measures ANOVA with two main effects (stimulus conditions) at a time.

To compare distributions of CPSD phase across stimulus conditions, we used the Matlab Circular Statistics Toolbox (Berens, 2009). We assessed CPSD phase angles for Intact, Measure, and Reversed responses only, since Phase stimuli did not contain a musical beat. For inter-subject analyses, we performed two-sided circular t-tests on the distribution of phase angles at the frequency of interest on a per-stimulus basis to determine whether each distribution differed significantly from a zero-degree-mean distribution. For stimulus-response analyses, we performed a circular ANOVA on a per-song basis to determine whether distributions of phase angles differed significantly by stimulus condition. Following this, we performed pairwise circular t-tests among pairs of stimulus conditions for each song to determine whether distributions of cross spectrum phase angles differed significantly by stimulus condition.

We corrected for multiple comparisons using False Discovery Rate (FDR) (Benjamini & Yekutieli, 2001), computing FDR-adjusted *p*-values in `R`. We report adjusted *p*_FDR_-values for analyses involving multiple comparisons.

### Data and Code Availability Statement

The raw EEG, preprocessed EEG input to RCA, and RC1-5 data, along with behavioral ratings and anonymized participant identifiers, are available for download from the Stanford Digital Repository (Kaneshiro et al., 2016a).^5^ Illustrative code reproducing the main results of the paper, along with RC1 data, are made publicly available via GitHub.^6^

## Results

We recorded scalp EEG responses from *N* = 48 participants listening to four full-length ‘Bollywood’ songs, each prepared in four conditions: In original form (Intact); with its measures randomly rearranged across time (Measure); with the entire waveform reversed (Reversed); and with its phase spectrum randomized (Phase). A subset of *N* = 12 participants heard each of the 16 stimuli twice (*N* = 24 trials per stimulus). For each condition, we computed an optimized measure of the neural correlation evoked during listening by spatially filtering the EEG to maximize the temporal correlation among response records—the inter-subject correlation (ISC) (Dmochowski et al., 2012; Parra et al., 2018). Based upon previous findings suggesting that ISC indexes emotional arousal or engagement with narrative stimuli, we hypothesized that both behavioral measures of enjoyment and ISC would be highest for the Intact stimuli, which were intended for consumption and have been shown to successfully engage a large audience.

### Intact stimuli elicit highest behavioral ratings

Overall, behavioral ratings aligned with our expectations of highest ratings for Intact stimuli composed for the listening audience, and lowest ratings for Phase stimuli, which are perceived as modulated noise (Song 1 waveforms shown in Figure 1A; Songs 2–4 waveforms included in Supplementary Figure S1). As suggested by visual inspection of Figure 1B, repeated-measures ANOVA performed on a per-question basis indicated that ratings covaried with stimulus condition for all questions (*χ*^2^ (3) ≥ 186.42, *p*_FDR_ < 2.2e-16, 4 comparisons). Follow-up pairwise analyses showed that Intact stimuli drew the highest ratings across songs for all four questions (*χ*^2^ (1) ≥ 27.91, *p_FDR_* < 1.6e-07, FDR corrected, 24 comparisons), while Phase stimuli were rated lowest for all questions (*χ*^2^ (1) ≥ 10.36, *p*_FDR_ < 0.0016). Ratings of Measure stimuli were found to be significantly higher than Reversed along dimensions of Pleasantness (*χ*^2^ (1) = 5.00, *p*_FDR_ = 0.028), Order (*χ*^2^ (1) = 5.11, *p*_FDR_ = 0.027), and Interest (*χ*^2^ (1) = 4.30, *p*_FDR_ = 0.040) but not Musicality (*χ*^2^ (1) = 2.72, *p*_FDR_ = 0.10).

### Stimuli retaining musical features produce consistent EEG components

Projected spatial filter weights can be visualized as topographic scalp maps. RC1 topographies, shown in Figure 2A, were virtually identical for Intact, Measure, and Reversed stimuli (all pairwise correlations *r* > 0.99). These topographies are similar to RC1 topographies reported by Cohen & Parra (2016) and Ki et al. (2016) in response to non-musical natural audio stimuli, and by Madsen et al. (2019)in response to natural music stimuli. They are also similar to component topographies derived from EEG responses to natural music using other spatial filtering techniques such as PCA (Schaefer et al., 2011, 2013), MUSIC(Sturm et al., 2015), and Canonical Correlation Analysis (CCA) (Gang et al., 2017). The spatial distribution of neural activity during Phase stimuli was markedly different, however (all pairwise correlations *r* < 0.73), hinting at a different neural generator for musical versus non-musical stimuli. In addition, as shown in Figure 2B, the RC1 correlation coefficient was well above permutation test significance thresholds for responses to Intact, Measure, and Reversed stimuli (all *p*_FDR_ < 0.0015, FDR corrected, 20 comparisons), but was not significant for the Phase condition (*p*_FDR_ = 0.070). RC correlation coefficients for the three musical stimulus conditions were markedly lower for components 2-5. Topographic maps for all computed RCs 1–5 are provided in Supplementary Figure S2.

**Figure 2:**
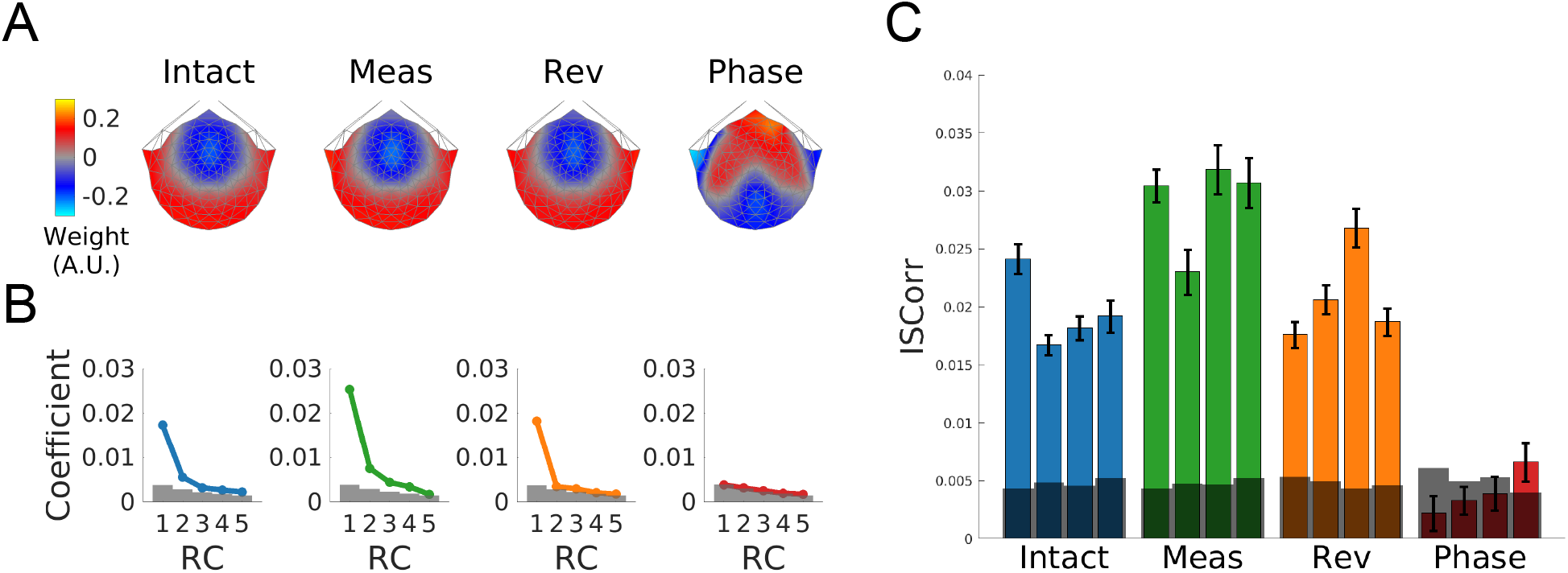
Measure-shuffled stimuli evoked highest inter-subject correlation (ISC). (**A**) Projected RC1 topographies were highly correlated across all stimulus conditions (all r > 0.99) except for Phase (all *r* < 0.73). (**B**) Correlation coefficients of RC1–5 for each stimulus condition. The shaded gray area denotes the 95th percentile of the null distribution. RC1 coefficients for Intact, Measure, and Reversed conditions were statistically significant (permutation test *p* < 0.01, FDR corrected) but dropped steeply after the first component; RC1 was not statistically significant for responses to Phase stimuli. (C) RC1 ISC was significant for all songs in Intact, Measure, and Reversed conditions (permutation test *p* < 0.01, FDR corrected) but not for Phase. ISC varied significantly by stimulus condition (repeated-measures ANOVA, *p* < 0.001), with highest ISC for Measure stimuli and lowest ISC for Phase (repeated-measures ANOVA, *p* < 0.001, FDR corrected). The shaded gray area denotes the 95th percentile of the null distribution; error bar height represents twice the standard error of the mean (*N* = 24 trials per stimulus).

### Measure stimuli evoke highest ISC

Because RC coefficients dropped steeply after the first component (Figure 2B), we focused our analyses solely on maximally correlated component RC1. We expected that neural correlation measured with ISC would mirror the trends observed in behavioral ratings. ISC was computed in a one-against-all fashion on RC1 EEG activations on a per-stimulus basis (24 trials per stimulus). As shown in Figure 2C, ISC was significant (all permutation test *p*_FDR_ = 0.0012, FDR corrected, 16 comparisons) across all songs for every stimulus condition except Phase. A repeated-measures ANOVA indicated that ISC differed significantly according to stimulus condition (*χ*^2^ (3) = 382.34, *p* < 2.2e-16). However, contrary to our expectation, whereas Intact stimuli always received the highest behavioral ratings, the most-correlated neural responses were evoked by the Measure stimuli: Follow-up comparisons among pairs of stimulus conditions revealed that ISC was significantly higher during Measure stimuli compared to all other conditions (*χ*^2^ (1) ≥ 72.01, *p*_FDR_ < 2.6e-16, FDR corrected, 6 comparisons). Responses to Phase stimuli had lowest ISC (*χ*^2^ (1) ≥ 165.29, *p*_FDR_ < 2.6e-16), and responses to Reversed stimuli had higher ISC than responses to Intact stimuli (*χ*^2^ (1) = 4.11, *p*_FDR_ = 0.043).

Our main findings of highest ISC for Measure stimuli and lowest ISC for Phase stimuli hold for summed ISC across RC1–3, as performed by Madsen et al. (2019); more information on summed RC1–3 coefficients and ISC is provided in Supplementary Figure S3. In addition, relating ISC to corresponding stimulus envelope fluctuations revealed negative correlations for Intact, Measure, and Reversed conditions, and a positive correlation for Phase (Supplementary Figure S4). All correlations were weak (|*r*| ≤ 0.15). Therefore, it is unlikely that the heightened ISC evoked by Measure stimuli was driven by magnitude envelope fluctuations.

### ISC correlates with envelope SRC

We sought to determine whether the measured EEG responses similarly tracked changes in stimulus envelopes and, if so, to what extent envelope following explained differences in ISC. To that end, we measured correlations between envelope fluctuations and RC1 EEG—the stimulus-response correlation (SRC). A similar approach has been used to decode brain states from single-trial EEG while experiencing natural audiovisual stimuli (Dmochowski et al., 2018) and intact music (Gang et al., 2017).

SRC results (Figure 3A) were similar to ISC results. First, SRC for each stimulus was significant (all permutation test *p* = 0.0012, FDR corrected, 16 comparisons) across all songs in every stimulus condition except Phase. Similar to the ISC case, a repeated-measures ANOVA revealed that SRC differed significantly according to stimulus condition (*χ*^2^ (3) = 178.13, *p* < 2.2e-16). Moreover, follow-up pairwise comparisons indicated that Measure stimuli produced highest SRC (*χ*^2^ (3) ≥ 18.95, *p*_FDR_ ≤ 1.3e-05, FDR corrected, 6 comparisons), and Phase stimuli produced lowest SRC (*χ*^2^ (3) ≥ 69.51, *p*_FDR_ < 4.4e-16). SRC did not differ significantly between Intact and Reversed stimuli. As ISC and SRC results were broadly similar, we computed their correlation on a per-trial basis; with 24 trials per stimulus and 16 stimuli, this procedure involved 384 trials. We observed a significant correlation (Figure 3B), with SRC explaining roughly 45 percent of the variance of ISC (*r* = 0.76, *p* = 1.01e-51).

**Figure 3:**
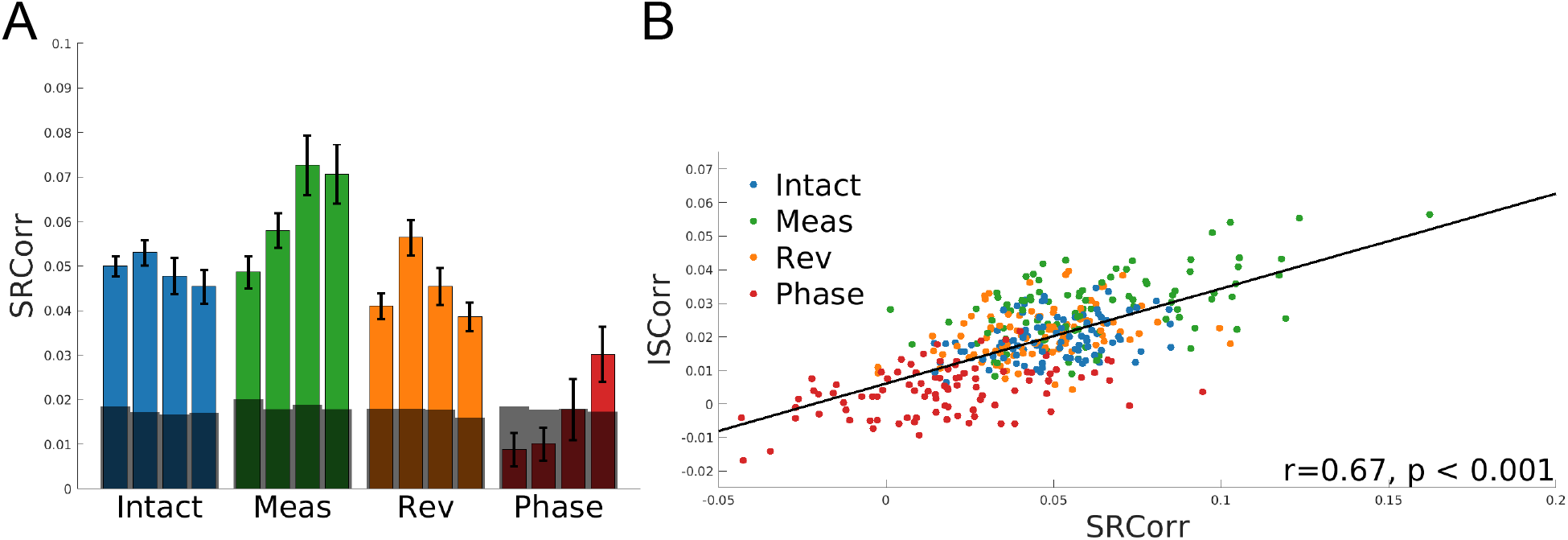
Stimulus-response correlation was highest for measure-shuffled stimuli. (**A**) Stimulus-response correlation (SRC), measuring the extent to which RC1 activations track fluctuations in stimulus envelope, was highest in the Measure condition, followed by Intact and Reversed, and lowest for Phase. All stimuli except Phase songs 1–3, exceeding the shaded gray area, produced statistically significant SRC (permutation test *p* < 0.01, FDR corrected). RC1 SRC varied significantly by stimulus condition (repeated-measures ANOVA *p* < 0.001). Follow-up pairwise comparisons showed that SRC was highest for Measure stimuli and lowest for Phase stimuli (repeated-measures ANOVA, *p* < 0.001, FDR corrected). (**B**) SRC and ISC of corresponding trials were significantly correlated, with SRC explaining around 45% of the variance of the ISC (*r* = 0.67).

### Frequency-domain EEG correlations implicate musical beat

To determine whether correlated temporal activations implicated perceptually relevant frequencies, we computed the magnitude-squared coherence spectrum (frequency-domain correlation) of RC1 activations for responses to Intact, Measure, and Reversed stimuli, which retained a steady beat. Coherence was computed among responses to a given stimulus (inter-subject) as well as between responses and envelope fluctuations (stimulus-response). We observed prominent peaks in the low-frequency coherence spectrum (0–12 Hz). As shown in Figure 4A, peaks occurred at frequencies corresponding to metrically relevant groupings and subdivisions of the musical beat for each song (e.g., at 5.2 Hz for Song 1, see Supplementary Figures S5–S7 for Songs 2–4). Interestingly, dominant peaks tended to occur in the 5–7 Hz range—a frequency often employed in studies of the visual system, but one which implicated different multiples of the perceptual tempo frequency (different beat ‘harmonics’) depending upon the tempo of each song.

**Figure 4:**
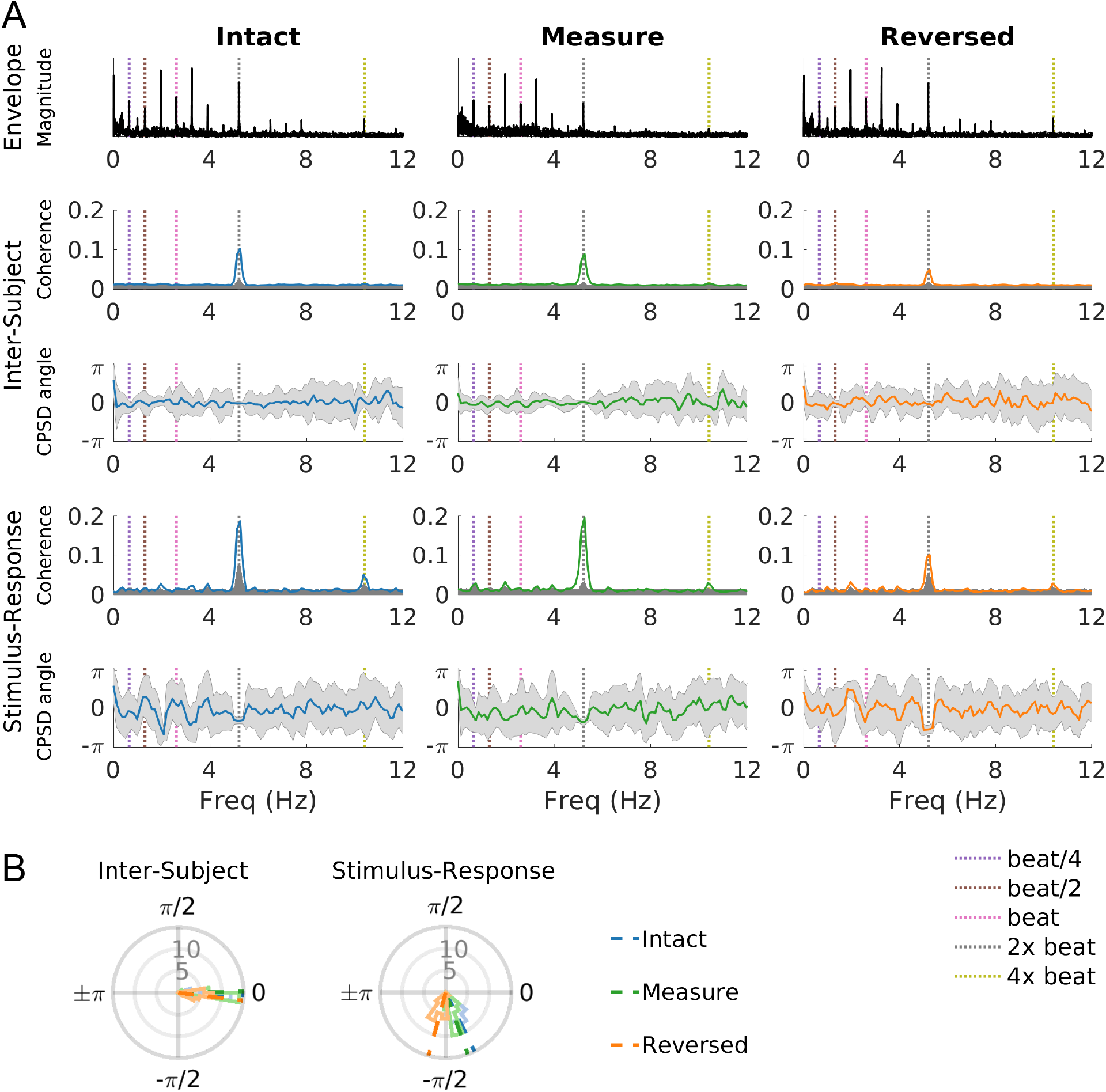
Song 1 coherence and cross power spectral density angles. (**A**) Envelope fluctuation spectra as well as coherence and CPSD angles for inter-subject and stimulus-response analyses at low frequencies. Prominent coherence peaks and convergence of CPSD phase angles were observed at frequencies related to the musical beat (here at 5.2 Hz). Coherence plots present the mean (colored line) and 95% significance threshold (shaded gray area). CPSD plots present the mean (black line) and standard deviation (shaded gray area). (**B**) Left: With one exception, distributions of inter-subject CPSD phase angles at the frequency of peak coherence for individual stimuli (across all songs) did not differ significantly from zero (circular t-test *p* > 0.05, FDR corrected). Right: Distributions of stimulus-response phase angles differed significantly according to stimulus condition for all songs (circular ANOVA *p* < 0.001, FDR corrected); distributions of angles for Reversed responses differed significantly from Intact and Measure (circular t-test *p* < 0.001, FDR corrected), while Intact and Measure never differed significantly from one another.

In a post-hoc analysis, we analyzed distributions of magnitude-squared coherence at the peak frequency of each stimulus (Figure 5A–B). Permutation testing revealed statistically significant coherence at the ‘frequency of interest’ for each stimulus in inter-subject (*p*_FDR_ < 0.001, FDR corrected, 12 comparisons) and stimulus-response (*p*_FDR_ < 0.001, FDR corrected, 12 comparisons) contexts. In addition, coherence distributions at these frequencies differed significantly according to stimulus condition (repeated-measures ANOVA, inter-subject *χ*^2^ (3) = 111.26, *p* < 2.2e-16; stimulus-response *χ*^2^ (3) = 41.239, *p* = 1.1e-09). However, follow-up pairwise comparisons did not align with time-domain correlation results: While the highest time-domain correlation was observed for responses to Measure stimuli, coherence did not differ significantly between Intact and Measure stimuli (inter-subject *χ*^2^ (1) = 1.13, *p*_FDR_ = 0.29; stimulus-response *χ*^2^ (1) = 0.058, *p*_FDR_ = 0.81, FDR corrected, 3 comparisons per analysis), although both conditions produced significantly higher peak coherence than Reversed stimuli (inter-subject *χ*^2^ (1) ≥ 76.75, *p*_FDR_ ≤ 3.3e-16, stimulus-response *χ*^2^ (1) ≥ 29.08 *p*_FDR_ ≤ 7.2e-08, FDR corrected, 3 comparisons per analysis). As was the case with time-domain correlation, inter-subject and stimulus-response coherence at frequencies of interest were highly correlated at the single-trial level, with stimulus-response coherence explaining 59 percent of the variance of the inter-subject coherence as shown in Figure 5C (*r* = 0.77, *p* = 1.08e-57). In sum, prominent beat-related coherence peaks were observed in the RC1 activations, but the level of coherence at these frequencies did not explain the differences in broadband neural correlation (ISC) or tracking of stimulus envelope fluctuations (SRC).

**Figure 5:**
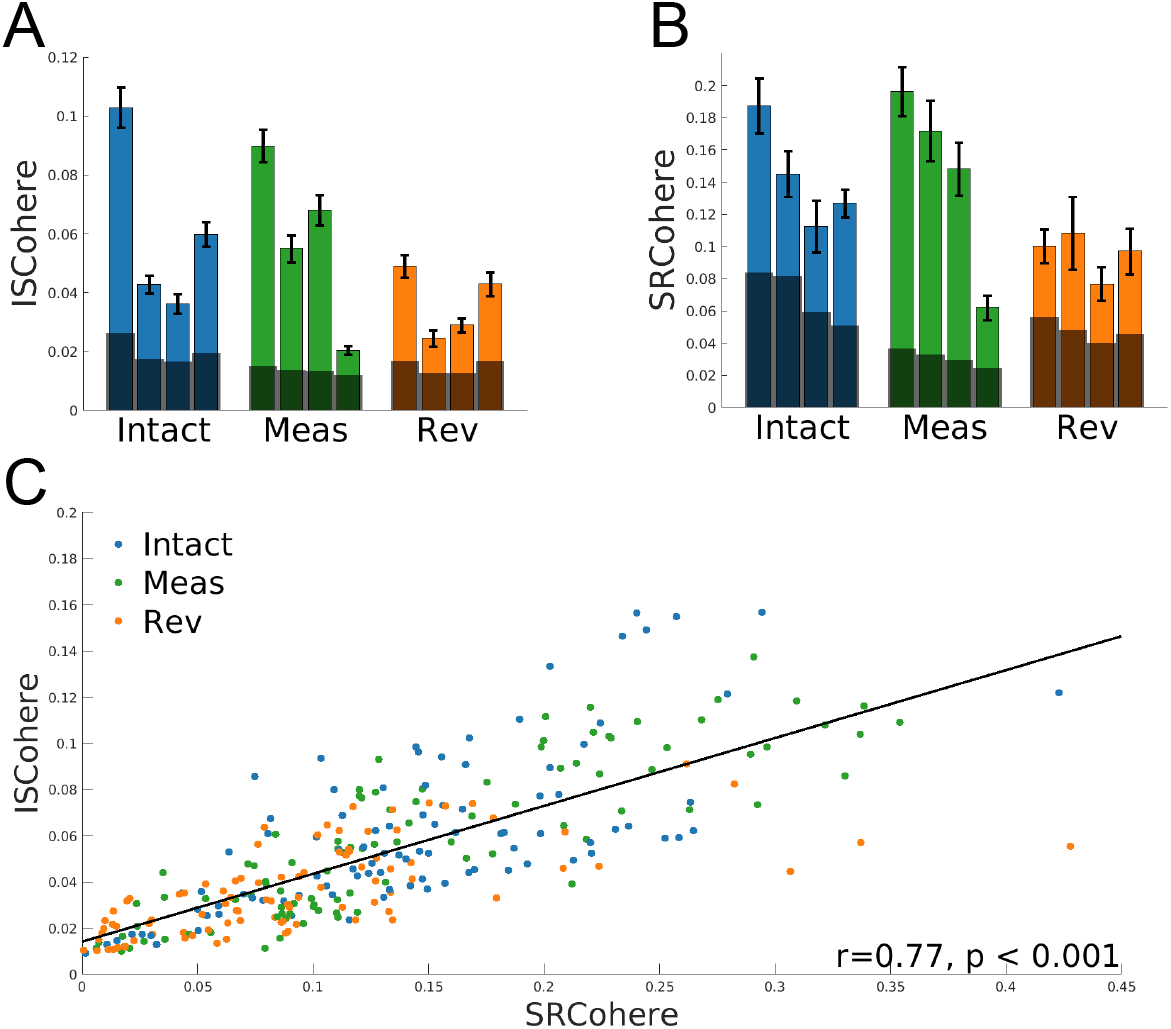
Peak coherence values. (**A**) Inter-subject coherence at each song’s frequency of interest was always statistically significant (permutation test *p* < 0.001, FDR corrected) and varied according to stimulus condition (repeated-measures ANOVA, *p* < 0.001). Intact and Measure coherence were higher than Reversed coherence (repeated-measures ANOVA, *p* < 0.001, FDR corrected) but did not differ significantly from one another. (B) Stimulus-response coherence at the same frequencies was also significant (permutation test *p* < 0.001, FDR corrected) and varied according to stimulus condition (repeated-measures ANOVA, *p* < 0.001), with higher coherence for Intact and Measure compared to Reversed (repeated-measures ANOVA, *p* < 0.001, FDR corrected). (C) Correlation of stimulus-response and inter-subject coherence for individual trials was significant at the peak frequency for the respective stimulus (*r* = 0.77), with stimulus-response coherence explaining 59% of the variance of inter-subject coherence.

### Stimulus reversal impacts cross power spectral density phase

RCA, the spatial filtering technique employed in this analysis, emphasized synchronous (i.e., in-phase) neural activity across responses to a given stimulus. However, maximizing correlation across EEG records does not elucidate stimulus-response time lags, and phase-locking would not necessarily occur at all frequencies. We accounted for the unknown lag during time-domain SRC analysis by constructing a temporal filter to account for all time sample-wise lags up to 1 second (see Methods; temporal filters are visualized in Supplementary Figure S8). In the frequency domain, we investigated lag by computing cross power spectral density (CPSD) phase angles among trials (inter-subject) and between each trial and corresponding stimulus envelope fluctuations (stimulus-response). Figure 4 indicates that responses were not phase-locked at all low frequencies. However, CPSD phase angles did tend to converge at frequencies corresponding to maximal coherence peaks (though this was not always the case; see Song 2 and Song 3 Reversed responses, Supplementary Figure S5–S6).

We next compared the distributions of phase angles at the frequency of interest implicated by coherence peaks by visualizing them as polar histograms (visualized for Song 1 in Figure 4B; results for other songs are shown in Supplementary Figure S5–S7). In the intersubject case, phase-locking would imply a zero-mean distribution of CPSD phase angles at this frequency regardless of stimulus tempo or condition. This proved to be the case: Circular t-tests on a per-stimulus basis indicated that with the exception of only one Reversed stimulus (Song 1), distributions of phase angles did not differ significantly from a zero-mean distribution (*p*_FDR_ > 0.05, FDR corrected, 12 comparisons).

Analysis of stimulus-response CPSD phase angles was performed on a per-stimulus basis in order to account for possible effects of both stimulus condition and tempo. Circular ANOVA results comparing phase angles across conditions of a given song were statistically significant for all songs (*p*_FDR_ ≤ 3.2e-12, FDR corrected, 4 comparisons), suggesting that distributions of stimulus-response phase angles did vary according to stimulus condition (Figure 4B). Follow-up pairwise comparisons revealed that for every song, distributions of Intact and Measure phase angles differed significantly from Reversed (*p*_FDR_ ≤ 3.3e-08, FDR corrected, 12 comparisons), but not from one another. Therefore, the stimulus-response lag at perceptually relevant frequencies was not modulated by disruption of short-term musical structure as in the Measure case, but was modulated by stimulus reversal.

## Discussion

ISC and SRC approaches have proven useful for studying neural processing of natural stimuli. In recent studies, temporal disruptions to such stimuli have been employed to disassociate low-level processing of stimulus features from processing of higher-level structure in audio and audio-visual narratives as well as music (Hasson et al., 2008; Lerner et al., 2011; Dmochowski et al., 2012; Regev et al., 2013; Farbood et al., 2015; Iotzov et al., 2017). In the present study we have applied this technique to natural music in order to examine the impact of temporal organization on neural correlation and listener enjoyment.

We presented participants with popular yet novel musical works in original form as well as in states of temporal disruption. We found that stimuli retaining basic musical features produced statistically significant ISC and SRC, while Phase excerpts did not (Figure 2-3). These results confirm findings of Abrams et al. (2013)—whose fMRI study also involved a phase-scrambled control—in that statistically significant ISC was not elicited by low-level auditory cues alone. Our results also extend their findings, as we find that natural music excerpts need not be in their original, intact form in order to elicit significant ISC. However, we note that among the musical features removed by phase scrambling are amplitude envelope fluctuations characteristic to music (Figure 1A). Therefore, subsequent investigation is aimed to disentangle the contributions of amplitude envelope fluctuations and higher-level acoustical and musical features in driving temporally correlated responses.

Our results also indicated that brain responses to synthetic music constructed in a way that produced a consistently high level of surprise (Measure condition) were most correlated in both ISC and SRC contexts. This finding was in contrast to our expectation, which was that Intact stimuli created for public consumption would produce the most correlated brain activity. These findings could not be explained simply by temporal discontinuities in stimulus envelopes, nor by coherence measures at beat-related frequencies. As expected, behavioral ratings of Pleasantness and level of Interest were highest for Intact versions of songs.

Our combined findings diverge from certain previous reports linking increased neural correlation to temporal intactness and to preference. For example, Farbood et al. (2015), using fMRI ISC, identified brain regions that were preferentially activated for intact musical stimuli compared to those scrambled at the measure level. Other previous EEG research has shown that scrambling an audiovisual film stimulus at the scene level—a manipulation that provided the impetus for our own Measure condition, as similar timescales are implicated— caused ISC to drop significantly (Dmochowski et al., 2012). A link between EEG ISC and participants’ preference ratings has also been established during viewing of intact television commercials (Dmochowski et al., 2014).

However, heightened temporal correlation of EEG has not always been reflective of intact or even comprehensible acoustical events. Ki et al. (2016) reported that when fluent English speakers were presented with an intact narrative, a narrative scrambled at the word level, and a narrative in a foreign language, ISC of RC1 EEG was highest for the foreign-language narrative and lowest for the intact English narrative. Factors other than enjoyment and preference are also known to modulate temporal correlation of EEG: Both ISC and SRC have been found to increase under heightened attention (e.g., with a more challenging task) and motor engagement, and to decrease when participants are distracted (Ki et al., 2016;Dmochowski et al., 2018; Ki et al., 2019).

Although the present disassociation between neural correlation and behavioral ratings of Pleasantness and Interest may suggest that neural correlation is not an effective index of musical engagement, an alternative interpretation is that the preservation of metrical structures, compounded by a moderate degree of expectation violation resulting from the disjunctions of the shuffled measures, are complicit in the higher correlations. Music is primarily non-representational; it is characterized by repetition and intentional disjunction, and is reliant upon structural hierarchies—both rhythmic and harmonic—nested within a metrical framework. Music listening is as reliant upon retrospection as it is upon prospection, in that non-lexical temporal units work in conjunction (and sometimes in opposition) to the metrical framework. Therefore, engagement here may point to a heightened state of attention associated with anticipating and processing continually surprising musical events of the Measure condition, even if listeners do not equate this state with enjoyment.

### Relating ISC and SRC

We studied neural correlation in time and frequency domains, both among EEG trials and between trials and stimulus envelope fluctuations. In both domains we found stimulus-response and inter-subject measures to be significantly correlated with one another. This finding aligns with past reports of significant EEG ISC-SRC correlation (Dmochowski et al., 2018). Our present component topographies and SRC results also align closely with those reported in a previous SRC study using the same Intact responses analyzed here, but which maximized SRC by jointly computing a spatial EEG filter and temporal stimulus filter using CCA (Gang et al., 2017). The correspondence between ISC and SRC is a reasonable finding since correlated responses are assumed to reflect exogenous (stimulus-evoked) processing (Dmochowski et al., 2012); in other words, maximally correlated activations should reflect some attribute(s) of the stimulus. We also found that SRC values tended to exceed ISC values, which confirms past findings (Dmochowski et al., 2018) and is also a reasonable result since ISC involves correlations among relatively noisy EEG responses, while SRC correlates EEG records to a cleaner stimulus feature.

### Complementary insights from ISC and SRC

Neurophysiological responses have been shown to track the envelope of speech (Aiken & Picton, 2008; Lalor & Foxe, 2010; Ding & Simon, 2012). While tracking of stimulus envelopes has been a popular approach in recent EEG work (O’Sullivan et al., 2015), it should be noted that stimulus-response approaches narrow the focus of analysis to individual features. In contrast, inter-subject analyses reflect holistic processing not requiring preselection of a stimulus feature, and it is unlikely that correlation as measured by inter-subject approaches would be fully explained by a single stimulus feature. However, our stimulus-response analyses have provided useful insights into the time course of EEG activations relative to time-varying stimulus features—for example, differences in temporal filters and CPSD phase angles for responses to Reversed stimuli—which could not have been attained from inter-subject analyses alone. Future work can explore this further using more holistic stimulus representations such as spectrograms, or by operating directly on the audio waveform.

### Evidence of Beat Processing

Periodicities of auditory and visual stimuli are known to elicit narrow-band evoked responses at frequencies directly related to those present in the input. In the case of music, such periodicities may include temporal regularities related to beat and meter. Spectra of EEG responses to rhythmic stimuli have been analyzed using a ‘frequency tagging’ approach, whereby trial-averaged responses are transformed to the frequency domain and spectral amplitudes at frequencies related to the musical beat are compared (Nozaradan et al., 2011, 2012). Magnitudes of responses at beat-related frequencies have been shown to exceed those at unrelated frequencies, even when relevant frequencies were endogenously imposed by the listener (as in imagining a duple or triple meter over a beat sequence, Nozaradan et al. (2011)) or were not dominant in stimulus envelope spectra (Nozaradan et al., 2012). We have extended this line of research using natural stimuli, taking an alternative analysis approach by computing RC1 frequency-domain correlation (coherence) among single-trial responses and between trials and corresponding stimulus envelopes.

We found significantly correlated activity at beat-related frequencies in both inter-subject and stimulus-response contexts. Largest peaks tended to occur at the tempo harmonic in the 5–7 Hz range regardless of stimulus tempo. For Song 1 this implied twice the tempo frequency, while for the other songs it implied four times the tempo frequency. As can be seen in Figure 4 and Supplementary Figure S5–S7, coherence peaks did not simply reflect frequencies of dominant spectral peaks of stimulus envelope fluctuations. Other stimulus-response coherence peaks implicated additional (sub-)divisions of the beat or musically relevant beat structures present in the songs. Notably, smaller coherence peaks at 3/4 the tempo frequency in Song 1 (Figure 4) and 3/2 the tempo frequency in Song 3 (Supplementary Figure S6) reflect complex rhythmic patterns found in these songs. The correlation of inter-subject and stimulus-response measures was higher in the frequency than time domain. This may be due to the fact that, given significant activations in narrow frequency bands corresponding to coherence peaks, we measured this activity with high SNR compared to broadband activations, as the noise bandwidth affecting the measurement was very small.

### Methodological contributions

Music is unique among natural stimuli in its capability to provide a hierarchically structured temporal framework through which periodic activations can be observed. Previous music research employing frequency tagging has typically operated on across-electrode averages of FFT magnitudes of response data, which themselves had already undergone trial averaging in the time domain. While averaging has been justified as a means of improving SNR (Nozaradan et al., 2012), our combined spatial filtering and coherence approach enabled us to identify correlated frequency-domain activations on the level of single trials of continuous, unepoched data. Trial averaging has also been reasoned to attenuate non-phase-locked activity, although it has been argued that magnitude-only measurements in the frequency domain may ignore important phase information (Rajendran & Schnupp, 2019). In our present analysis, the weighted spatial averaging of RCA—which maximized temporal covariance across trials—and the coherence measure emphasized phase-locked activity. Finally, we observed phase directly by computing CPSD phase angles; we found RC1 responses to be largely in phase at frequencies with significant coherence, and observed significant differences in phase lag for Reversed stimuli.

Finally, past research using frequency tagging to study beat perception has faced a substantial noise floor in the low-frequency EEG spectrum (Losorelli et al., 2017), which is typically corrected by manually ‘flattening’ the spectrum through subtraction of mean values from adjoining frequency bins (Nozaradan et al., 2011). The coherence approach used here removed the need to flatten the spectrum, and our permutation testing approach for statistical significance produced readily interpretable results. Further research using both coherence magnitude and CPSD phase can continue to bridge between controlled, synthesized stimuli and natural music to deepen our understanding of musical beat features and corresponding neural responses.

### Does neural correlation require a steady beat?

We intentionally chose stimuli containing steady, electronically produced beats and observed evidence of those temporal regularities in the coherence spectra and the temporal filters computed during SRC. However, peak coherence values did not explain the temporal correlation of the responses. We argue that this would not be expected to be the case, as music should not require a steady beat in order to evoke correlated responses. In fact, expressive variations in timing are known to constitute an important part of the music listening experience (Patel, 2010; Istók et al., 2013). This point has been verified by Madsen et al. (2019), who obtained significant EEG ISC using musical stimuli without steady, electronically produced beats. As such, while we observed correlated frequencies as a result of a steady beat, beat is not necessary (or perhaps even sufficient) to produce significant neural correlation. Future research can further explore the role of temporal regularities, and deviations from them, in eliciting correlated neural responses to natural music.

### Music-Specific Implications

Both this study and the recent study by Madsen et al. (2019) considered only aggregate measures of neural correlation—that is, correlations computed across the entire duration of a stimulus. However, temporally resolved EEG ISC has been shown to peak during periods of high tension and suspense in audiovisual film excerpts (Dmochowski et al., 2012). The additional response features offered by that approach—such as quantity, timing, height, and duration of significant correlation peaks—will enable neural measures to be interpreted in relation to specific occurrences of musical analogues of narrative devices, as has been suggested during periods of buildup to structural high points and at structural segmentation boundaries (Kaneshiro, 2016; Kaneshiro et al., 2016b).

We obtained significantly correlated responses to musical stimuli heard during a single listening session. Needless to say, a standard measure of the ‘success’ of a musical work is the music’s enticement to hear the work repeatedly. In fact, musical preference often develops over multiple hearings (Madison & Schiölde, 2017). While Madsen et al. (2019) reported sustained ISC across repeated hearings of unfamiliar music, future research can take a more nuanced approach to investigate the interplay of engagement, preference, multiple hearings, degree of focus, and modes and contexts of listening. A longitudinal approach would better characterize features of music that drive increasingly reliable responses over time. The unique attributes of music and practices of music consumption, as compared to other types of stimuli, promise such heightened nuance.

Finally, lyrics constitute an important, and for many listeners, integral component of the enjoyment of music. Numerous studies pursue evidence for the degree to which words and music are processed independently or integrated with one another (Gordon et al., 2010;Besson et al., 1998), while high-level musical descriptors such as genre (Fell & Sporleder, 2014;Tsaptsinos, 2017) and mood (Hu & Downie, 2010) have been predicted from lyrics alone. From an experimental design perspective, lyrics serve as a potential confound and complicate stimulus manipulations. Given that musical works consumed by typical demographics of participants (college undergraduate and graduate students) are songs with lyrics, this poses a challenge to studying musical engagement with truly ecologically valid stimuli. Studies typically avoid this confound by using instrumental music as stimuli (for example, Madsen et al. (2019)). As an alternative approach, here we have used popular music with lyrics in an unfamiliar language and obtained correlated responses even though the lyrics were incomprehensible to participants. Future research could focus specifically on lyrics in order to clarify their role in engaging listeners.

## Conclusion

Music is a pervasive human activity. Humans engage with music both passively and actively, both for its own pleasure and for a purpose. Music is employed functionally and therapeutically to regulate emotion (Moore, 2013). It can strengthen autobiographical memory (Irish et al., 2006), promote motor activity (Thaut & McIntosh, 2014), affect temporal perception (Kellaris & Kent, 1992; Droit-Volet et al., 2013), and foster social interaction and bonding (Tarr et al., 2014). Music’s therapeutic and rehabilitative powers (Thaut & McIntosh, 2014) have been consistently noted since the ancient Greeks (Provenza, 2019). Many of the behavioral states associated with or caused by music—including, among others, memory retention and temporal perception—are suggested by neural correlation during natural music listening. Documenting and measuring neural measures of engagement with ecologically valid stimuli as reported here promises deeper insights into musical engagement.

We have demonstrated that inter-subject and stimulus-response EEG correlation techniques can be employed in the time and frequency domains to study natural music processing. Our results suggest that neural correlation during music listening may be driven in part by expectation formulation (encountering and processing novel content) and expectation processing (particularly violated expectation), and that a state of listener engagement can be disassociated from behavioral reports of enjoyment. These findings enrich the interpretation of elevated neural correlation as engagement and compel future studies to clarify the relationship between neural correlation, engagement, and enjoyment.

## Acknowledgments

We gratefully acknowledge Jay Kadis and Vladimir Vildavski for help with hardware configurations, and Wenfei Du for statistical advice.

## Abbreviations

(CPSD): Cross power spectral density;
(FDR): False Discovery Rate;
(ISC): inter-subject correlation;
(RC1): Reliable Component 1;
(RCA): Reliable Components Analysis;
(SS-EP): steady-state evoked potential;
(SRC): stimulus-response correlation.

## Supplementary Figures

**Figure S1:**
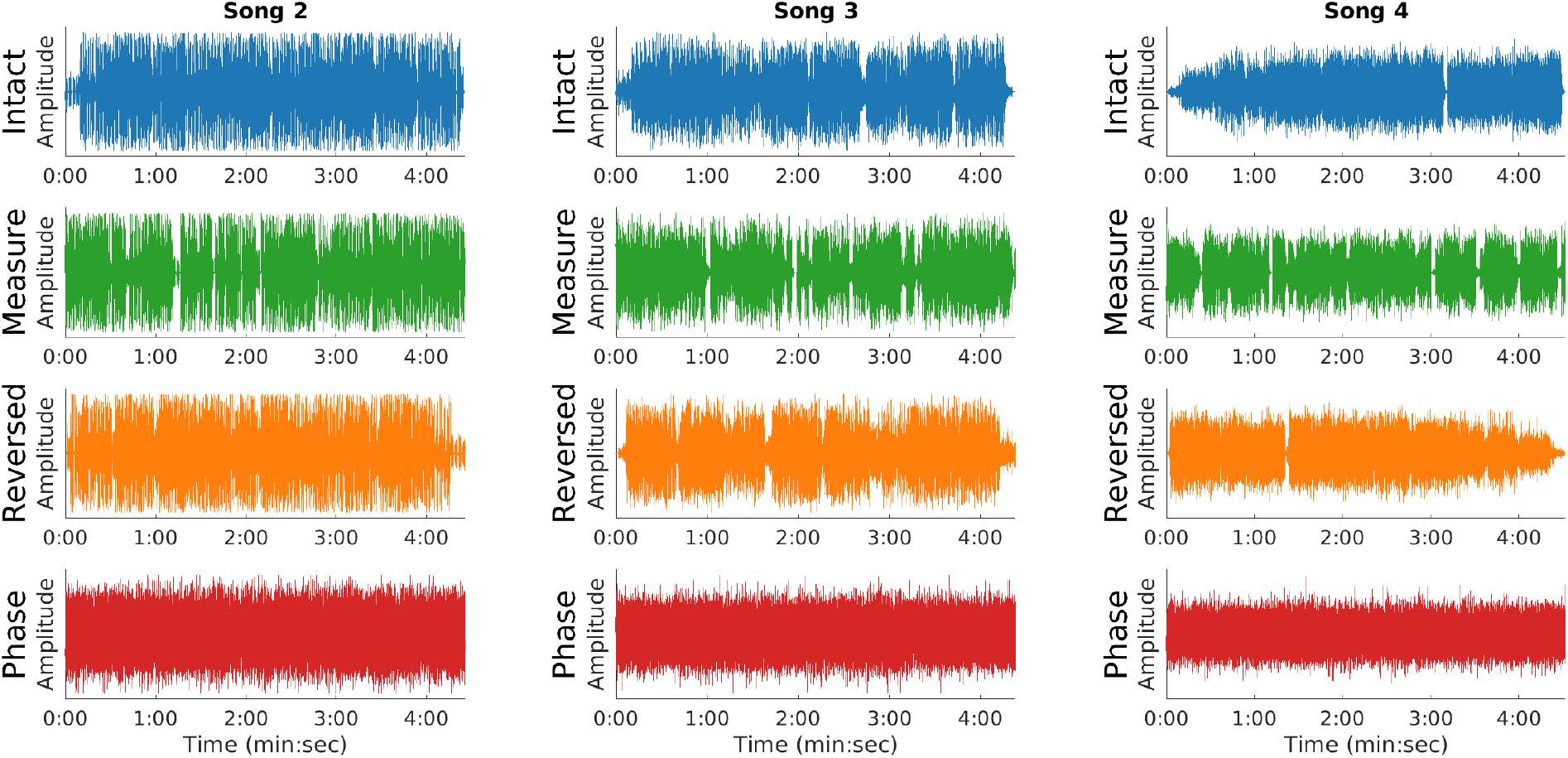
Stimulus waveforms for Songs 2–4.

**Figure S2:**
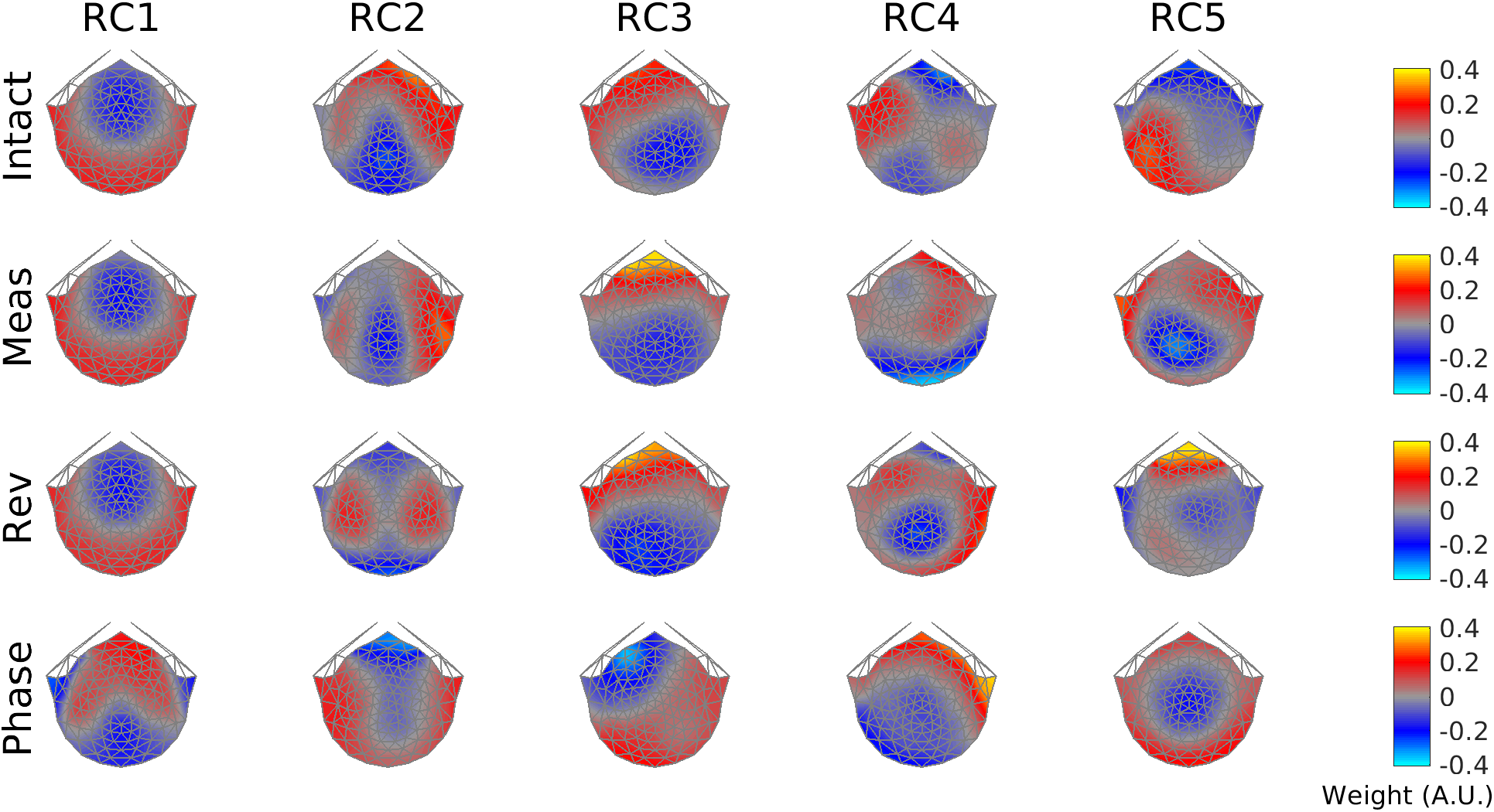
Forward-model projections of spatial filters, RCs 1–5.

**Figure S3:**
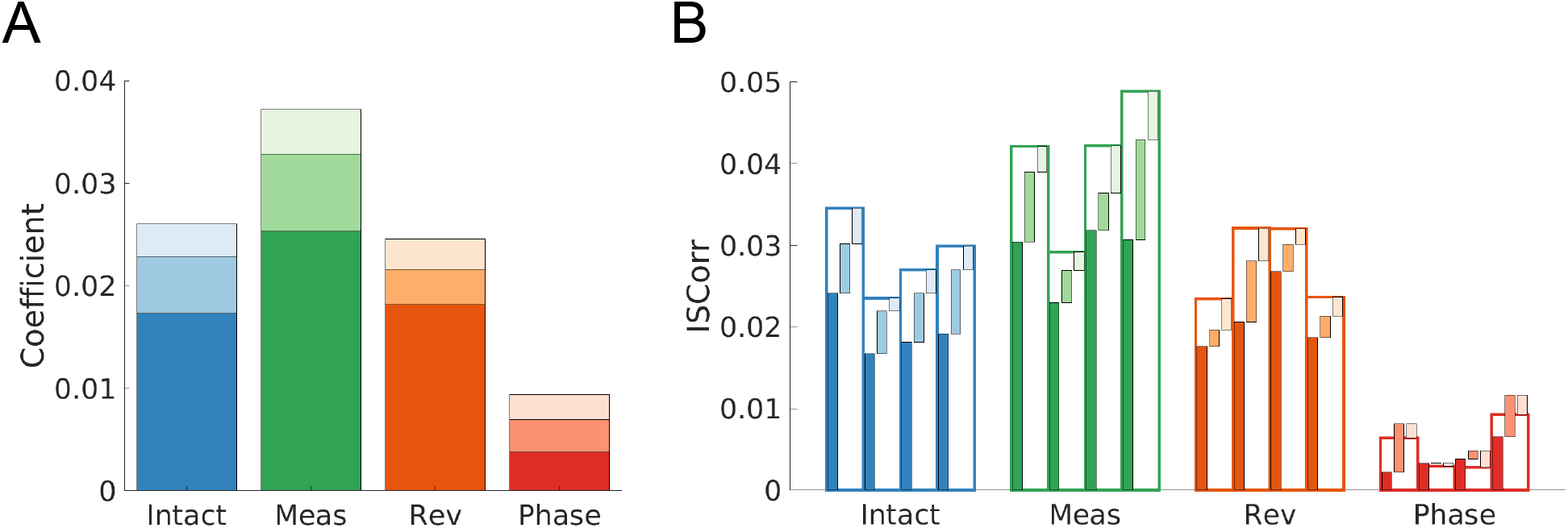
Summed correlation coefficients and ISC for RC1–3. (A) Summed correlation coefficients for RC1–3 were highest for responses to Measure stimuli and lowest for responses to Phase stimuli. (B) For each stimulus, ISC was computed separately for RCs 1–3 and then summed. A repeated-measures ANOVA indicated that summed ISC differed significantly according to stimulus condition (*χ*^2^ (3) = 305.83, *p* < 2.2e-16). Follow-up pairwise comparisons showed that Measure stimuli elicited highest summed ISC across RC1–3 (*χ*^2^ (1) ≥ 50.69, *p*_FDR_ ≤ 1.3e-12, FDR corrected, 6 comparisons), and Phase stimuli elicited lowest RC1–3 ISC (*χ*^2^ (1) ≥ 129.05, *p*_FDR_ ≤ 4.4e-16). In fact, ISC for Phase RCs 2 and 3 was sometimes negative.

**Figure S4:**
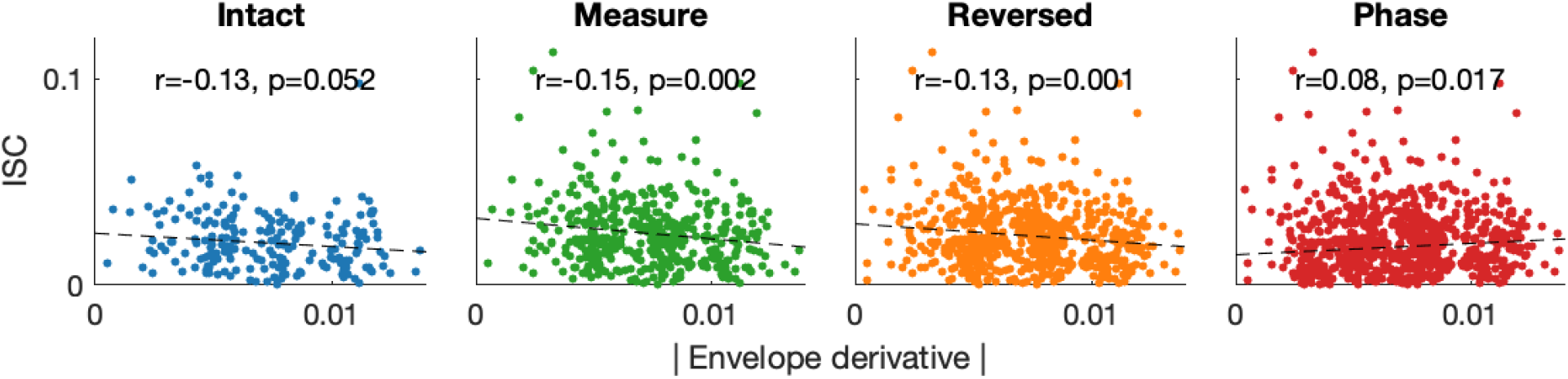
ISC is inversely related to envelope dynamics in musical stimuli. We computed both the ISC and the absolute mean of the envelope derivative along 5-sec time windows of all presented stimuli. Results were pooled across songs. For two of the three stimulus conditions retaining musical features, we found a significant inverse relationship between ISC and the amount of fluctuation in the envelope. For phase-scrambled stimuli, a small but significant positive correlation between ISC and envelope fluctuation was observed. Overall, all correlations were weak (|*r*| ≤ 0.15). These findings indicate that conditional differences in envelope dynamics do not explain those in the ISC (Figure 2C).

**Figure S5:**
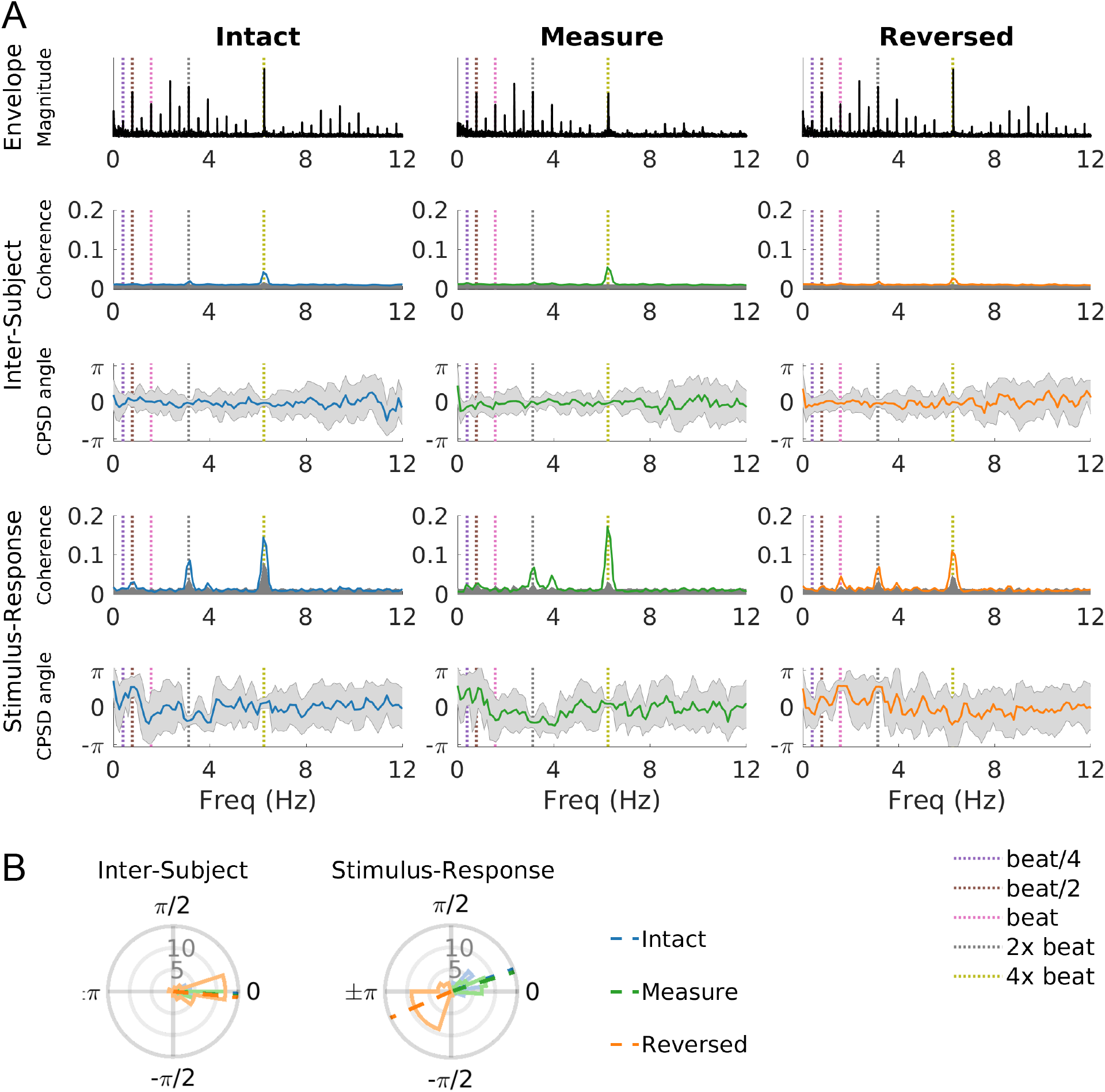
Coherence and cross power spectral density phase angles for Song 2. Peak coherence is observed at 6.3 Hz (four times the tempo frequency).

**Figure S6:**
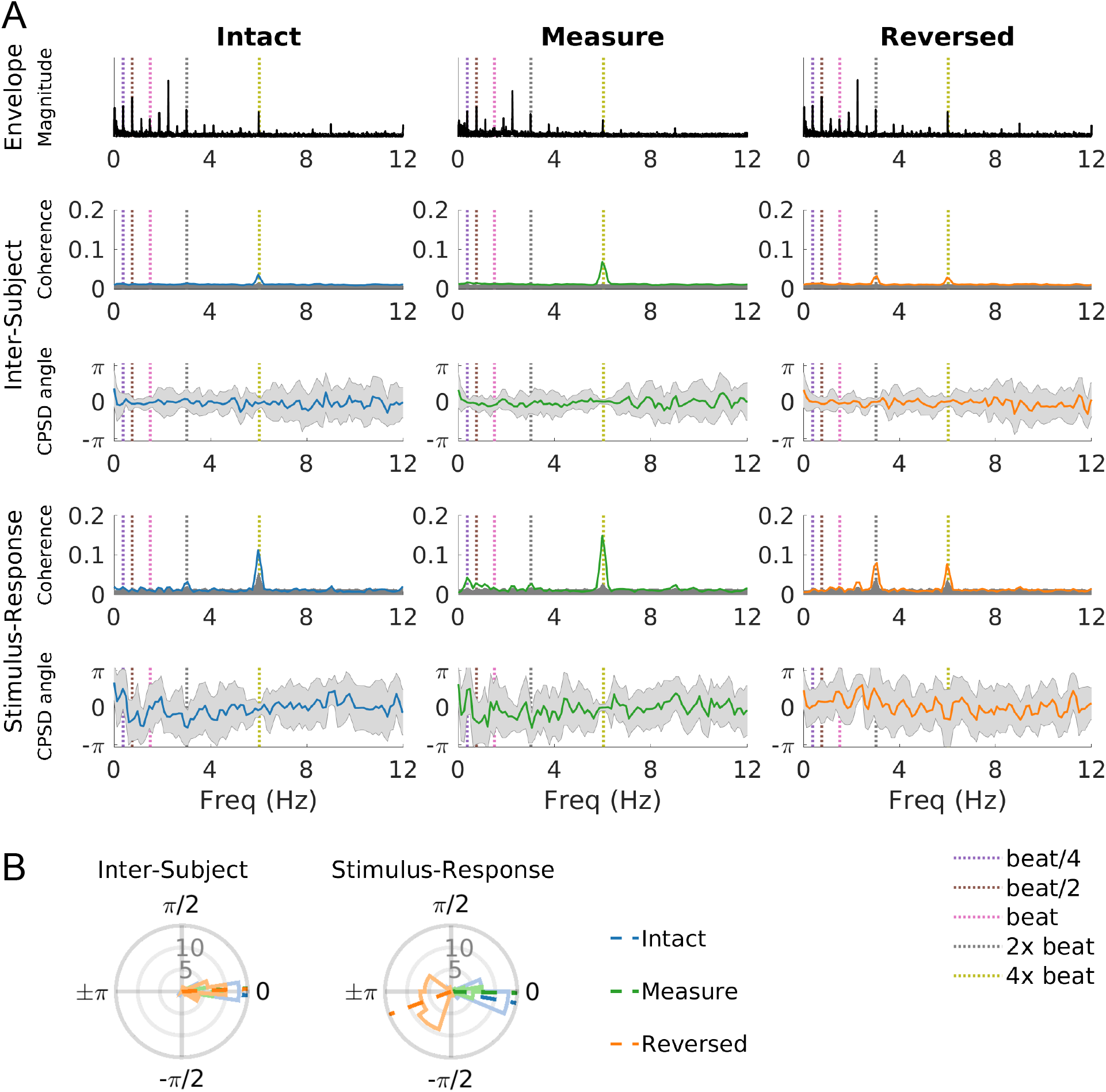
Coherence and cross power spectral density phase angles for Song 3. Peak coherence is observed at 6.0 Hz (four times the tempo frequency).

**Figure S7:**
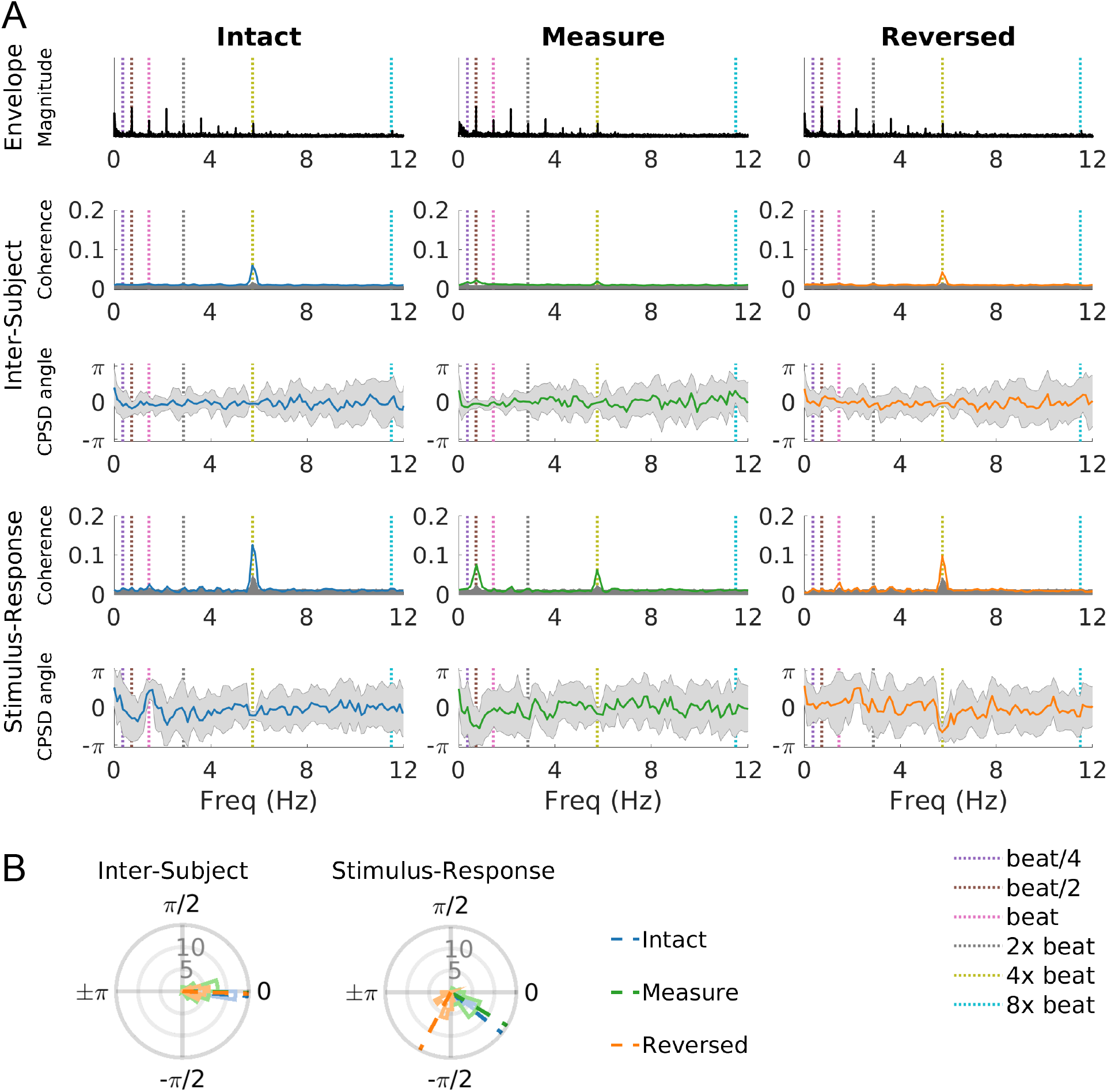
Coherence and cross power spectral density phase angles for Song 4. Peak coherence is observed at 5.7 Hz (four times the tempo frequency).

**Figure S8:**
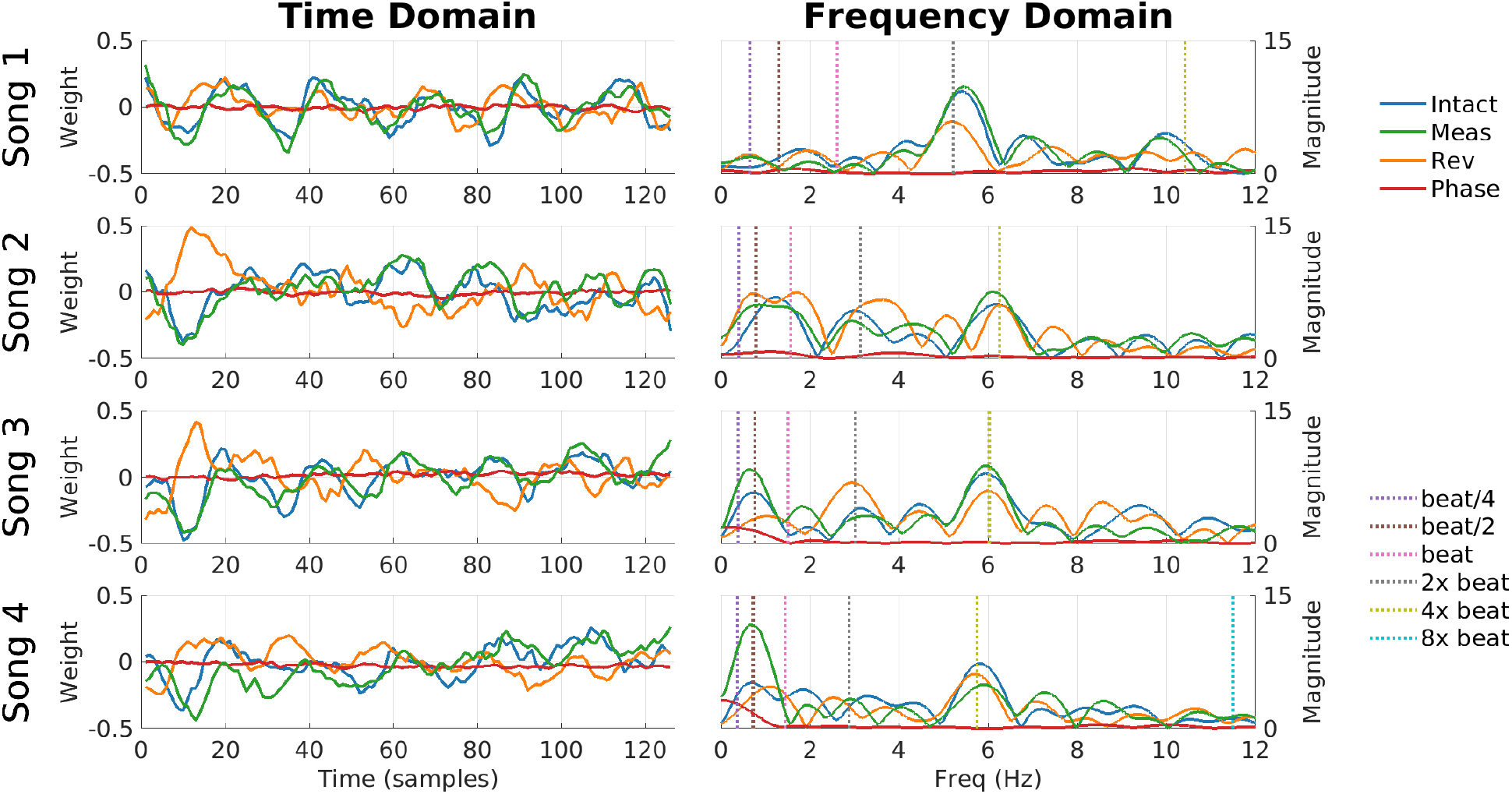
Temporal filters computed during stimulus-response correlation, visualized in the time and frequency domains.

## Footnotes

1 Audacity^®^ is copyright ©1999–2016 Audacity Team, http://audacity.sourceforge.net/

2 https://www.mathworks.com/

3 https://www.neurobs.com/

4 https://github.com/dmochow/rca

5 https://purl.stanford.edu/sd922db3535

6 https://github.com/blairkan/music-isc-illustrative

